# Kif2 microtubule depolymerase is required for unequal cell division and localizes to a subdomain of cortical endoplasmic reticulum

**DOI:** 10.1101/171041

**Authors:** Vlad Costache, Celine Hebras, Gerard Pruliere, Lydia Besnardeau, Margaux Failla, Richard R. Copley, David Burgess, Janet Chenevert, Alex McDougall

## Abstract

Unequal cell division (UCD) is a fundamental process responsible for creating sibling cell size asymmetry; however, how microtubules are specifically depolymerized on one aster of the mitotic spindle creating a smaller sibling cell during UCD has remained elusive. Using invertebrate chordate embryos (ascidian) that possess a large cortical structure (CAB) that causes UCD, we identified a microtubule depolymerase (Kif2) involved in creating cell size asymmetry. Kif2 localizes to the cortical subdomain of endoplasmic reticulum in the CAB. During three successive UCDs, Kif2 protein accumulates at the CAB during interphase and is delocalized from the CAB in mid mitosis. Rapid imaging of microtubule dynamics at the cortex revealed that microtubules do not penetrate the CAB during interphase. Inhibition of Kif2 function prevents the development of mitotic aster asymmetry and centrosome movement towards the CAB thereby blocking UCD, whereas locally increasing microtubule depolymerization causes exaggerated asymmetric spindle positioning. This study provides insights into the fundamental process of UCD and for the first time shows that a microtubule depolymerase is localized to a cortical site controlling UCD.

---

Control of microtubule dynamics at the cell cortex is important for a myriad of processes including spindle positioning at the cell center during cell division (Garzon-Coral et al., 2016; Kern et al., 2016; Minc and Piel, 2012), asymmetric spindle positioning during unequal cell division (UCD) in embryos (Cowan and Hyman, 2004; Kotak and Gönczy, 2013), and for axonal pruning during nervous system development in mammals (Homma et al., 2003; Maor-Nof et al., 2013). Microtubule dynamics have been intensively studied during asymmetric cell division (ACD) which is sometimes coupled with UCD creating one large and one small daughter cell that have different fates, as in *Drosophila* neuroblasts (Siller and Doe, 2009), sea urchin micromeres (Wessel et al., 2014) and *C. elegans* 1-cell zygotes (Galli and van den Heuvel, 2008). Such UCD relies on the development of unbalanced forces that has two components: cortical pulling force acting on astral microtubule plus ends (Grill et al., 2001) and depolymerization of microtubule plus ends as they encounter the cortex (Kozlowski et al., 2007; Labbé et al., 2003) which create unbalanced forces to position the mitotic spindle asymmetrically. These two processes acting together overcome the forces that cause mitotic spindles to move to the center of the cell, a process that senses and integrates force over the length of microtubules (Minc et al., 2011). However, one key piece missing from this model of UCD is the identity of the protein(s) that cause astral microtubule plus end depolymerization at the cortex, which is important not only for UCD but also for the mitotic spindle centering mechanism based on astral microtubule length that operates during symmetric cell division.

Microtubule plus ends can be induced to depolymerize via different mechanisms. *In vitro* experiments indicate that dynein can cause catastrophe of microtubule plus ends (Laan et al., 2012), raising the possibility that in intact cells dynein couples pulling with depolymerization. A different mechanism regulates microtubule plus end depolymerization in developing mammalian neurites which is dependent on the cortically-localized microtubule depolymerase Kif2A (Homma et al., 2003). Kif2A is a member of the kinesin-13 family of microtubule depolymerases (Lawrence et al., 2004) which includes MCAK/Kif2C that causes microtubule plus end depolymerization at kinetochores during anaphase (Wordeman et al., 2007). However, in cells that divide unequally it is still not known what causes astral microtubule plus end depolymerization at the cortex, and the only protein known to limit microtubule growth at the cortex during UCD in one cell *C.elegans* embryos does not affect astral microtubules (O’Rourke et al., 2010).

*C.elegans* embryos have provided a wealth of knowledge about the cortical pulling forces that act upon astral microtubules of the spindle during UCD. For example, following fertilization and symmetry breaking *in C. elegans*, the Par polarity complexes are partitioned to distinct cortical subdomains (Gönczy, 2008). Anterior Par3/Par6/aPKC (PKC-3) phosphorylates LIN-5 (NuMA) at the anterior cortex inhibiting the cortical anterior spindle pulling forces (Galli et al., 2011), while NuMAs binding partner GPR-1/2 (Pins/LGN) becomes enriched at the posterior cortex during mitosis (Gotta et al., 2003). The Dynactin/Dynein complex protein DRBY-1 interacts with Pins and NuMA thus providing a cortical link to microtubules (Couwenbergs et al., 2007). During metaphase and anaphase the mitotic spindle is pulled towards the posterior cortex causing UCD (Grill et al., 2001). Late in mitosis the posterior centrosome has changed from a spherical shape to a flattened and disc-shaped structure (Severson and Bowerman, 2003). Symmetric cell divisions in somatic cells also depend upon cortical dynein to center the mitotic spindle (Collins et al., 2012; Kern et al., 2016; Kiyomitsu and Cheeseman, 2013). In addition to dynein pulling forces, it has been shown that astral microtubule depolymerization also plays a role in posterior spindle displacement. For example, pulling forces are lacking when microtubules are stabilized by taxol, while in embryos carrying a temperature-sensitive mutation in a β-tubulin gene the posterior displacement distance of the spindle is enhanced (Nguyen-Ngoc et al., 2007). Based on these and other data a dual force-generation mechanism has been proposed that relies on microtubule pulling forces (dynein-dependent) combined with microtubule depolymerization (Kozlowski et al., 2007; Nguyen-Ngoc et al., 2007). Thus in *C.elegans,* although cortical microtubule depolymerization is thought to be part of the mechanism for posterior spindle displacement, the mechanism regulating cortical microtubule depolymerization is not known (Gönczy, 2008).

Many embryos provide more extreme examples of UCD whereby the two asters of the mitotic spindle become highly asymmetric in size and shape with the smaller of the two asters being inherited by the smaller of the two daughter cells. Similar to the flattened posterior centrosome in *C.elegans* one-cell embryos (Severson and Bowerman, 2003), such mitotic aster asymmetry has also been observed during UCD in spiralian (Lambert, 2010; Rabinowitz and Lambert, 2010), echinoderm (Holy and Schatten, 1991; Schroeder, 1987), and ascidian (Hibino et al., 1998; Prodon et al., 2010) embryos. Mitotic aster asymmetry commonly occurs at the third cleavage in spiralian embryos with the smaller aster associating with an animal cortical domain at the 4-cell stage leading to UCD (Rabinowitz and Lambert, 2010). A similar phenomenon has been documented for sea urchin embryos where one aster in each future micromere associates with a cortical domain situated at the vegetal pole of the 8-cell stage embryo creating aster asymmetry during UCD (Schroeder, 1987; Holy and Schatten, 1991). The clearest example of a cortical domain that causes aster asymmetry and UCD comes from ascidian embryos. Here a large cortical structure termed the CAB (centrosome-attracting body) causes three successive rounds of UCD accompanied by the CAB-proximal aster becoming smaller (Hibino et al., 1998; Nishikata et al., 1999; Prodon et al., 2010). Although the Par complex (aPKC, Par3 and Par6) is localized to the CAB (Patalano et al., 2006), we do not currently know how the CAB affects microtubule dynamics leading to aster asymmetry. Due the large dimensions of the CAB (circa 20µm at the 8-cell stage) we wondered whether it would be possible to identify proteins involved in regulating microtubule dynamics at the cortex via live cell imaging during UCD in the early ascidian embryo.

We have developed the optically transparent ascidian species *Phallusia mammillata* as a system to perform live cell imaging to study microtubule dynamics during UCD (McDougall et al., 2015; Prodon et al., 2010). By analyzing microtubule dynamics at the cortex we discovered that a microtubule depolymerase (Kif2) localizes to the cortical CAB in a cell-cycle dependent manner. Through live cell imaging of Kif2::Venus/mCherry/Tomato and immunofluorescence we demonstrate that exogenous and endogenous Kif2 localizes to the cortical CAB. In particular, Kif2 accumulates on a domain of cortical endoplasmic reticulum (cER) concentrated at the CAB during interphase and leaves the CAB cER during mid mitosis when CAB proximal microtubules become short. In addition, we show that microtubules do not penetrate the cortical CAB during interphase. Finally, we found that inhibiting endogenous Kif2 protein function prevents the establishment of mitotic aster asymmetry and UCD, and conversely that increasing depolymerization of microtubules near the CAB causes the CAB proximal spindle pole to move nearer to the CAB.

## RESULTS

### Unequal cell division and the CAB

The early embryo of the European ascidian *Phallusia mammillata* is favorable for live-cell imaging and functional studies because its cells are transparent (see Movie S1) and readily translate exogenous mRNAs such as those encoding GFP fusions and dominant negative constructs (McDougall et al., 2015). Ascidian embryos display three rounds of UCD that depend upon the centrosome-attracting body/CAB (Hibino et al., 1998; Nishikata et al., 1999; Prodon et al., 2010). The mitotic spindle aligns relative to the cellular polarity with one spindle pole pointed towards the CAB (Figure 1A). During these three rounds of UCD, astral microtubules emanating from the centrosome that polymerize towards the CAB and midline are shorter than those microtubules that do not polymerize towards the CAB or microtubules originating from the more distant centrosome as the mitotic spindle approaches the CAB and midline (Figure 1 A). A smaller and flattened aster thus forms nearest the CAB and midline during each round of UCD (Figure 1 and Movies S2, S3 and S4 with a 3D rendering of an 8-cell stage embryo shown in Movie S5). In *Phallusia* embryos both centrosomes appeared similar for γ-tubulin staining (sFig. 1) indicating that aster size is not a function of γ-tubulin loss from one centrosome as has been observed in leech zygotes (Ren and Weisblat, 2006).

**Figure 1.**
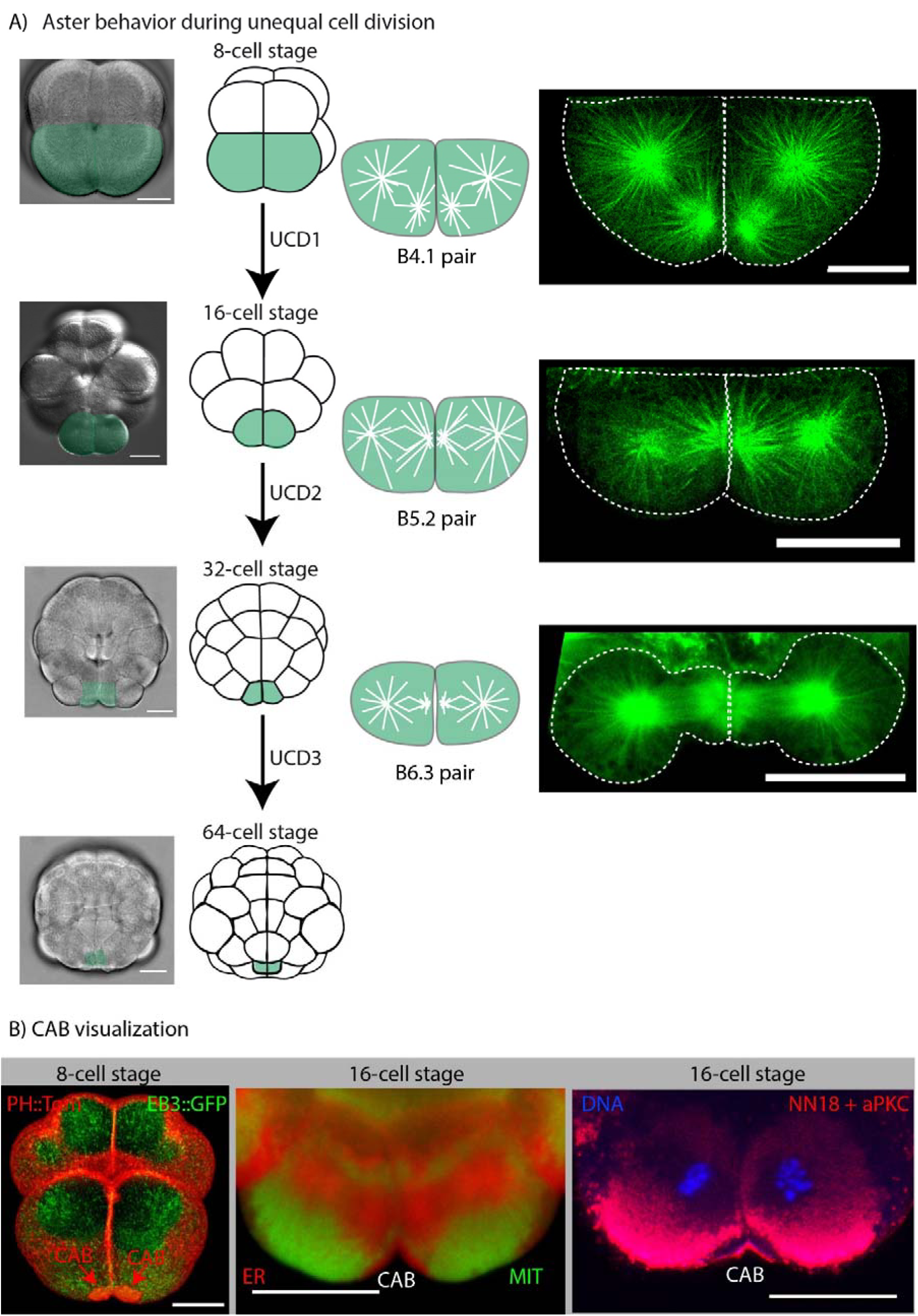
Microtubules and centrosome-attracting body (CAB) during UCD. (A) Aster behavior during unequal cell division. Schematics showing embryos from the 8 to 64-cell stage together with bright-field images of embryos with the blastomeres that undergo unequal cell division highlighted green. Right: selected confocal planes from live 4D imaging experiments showing microtubule organization in the pairs of blastomeres that undergo UCD at the 8, 16 and 32-cell stages (corresponding to the schematics). All microtubules were labelled in live embryos with Ensconsin::3GFP. Scale bars = 30µm. See Movies S1, S2, S3 and S4. (B) CAB visualization. Left image: 3D rendering of several confocal imaging planes reveals the CAB in the two bottom blastomeres (B4.1 pair) at the 8-cell stage in a live embryo (red arrows). CAB visualized with PH::Tom (red) and the deeper cytoplasm with the microtubule-binding protein EB3::GFP (green). See Movie S6. Center image: Live confocal imaging of the cortical endoplasmic reticulum in the CAB (labelled red) with a marker for endoplasmic reticulum (DiIC16) and the mitochondria (labelled green) with Mitotracker. The cER present in the CAB (also see deeper ER accumulated on the mitotic spindles) is attached to a specialized apical domain. Right image: Confocal image of a fixed 16-cell stage embryo stained with antibodies to cortical aPKC and mitochondrial ATP synthase showing the apical sub-domain lying at the cortical surface of the CAB and the deeper and surrounding mitochondria with NN18 respectively. Chromosomes stained with DAPI (blue). Note the dark unlabeled zone between the CAB surface and the mitochondria which is filled with cER. Scale bars = 30µm.

The CAB is a multilayer structure and can be visualized in several ways (Paix et al., 2011a; Patalano et al., 2006). Because the CAB creates a protrusion it can be visualized with plasma membrane markers such as PH::Tom (Figure 1 B and Movie S6). Also, because the CAB is rich in cortical ER and excludes mitochondria it can be visualized by specific lipophilic dyes that label either the mitochondria or cER in the CAB (Figure 1 B). Finally, antibodies to aPKC label the cortical surface of the CAB but do not label the cER which appears as a dark zone surrounded by mitochondria labelled with anti mitochondrial antibody-NN18 (Figure 1 B).

### Characterization of the microtubule depolymerase Kif2 in ascidian embryos

In order to understand how the CAB may be involved in creating aster asymmetry we searched for CAB-resident proteins by screening likely candidates by either probing with antibody for immunofluorescence or localization of expressed fluorescently tagged proteins. We identified a number of proteins including Kif2, a member of the kinesin-13 family of proteins that instead of possessing motor activity displays microtubule depolymerization activity (Hirokawa and Takemura, 2004). Vertebrates contain three members of the Kif2/kinesin-13 family: Kif2a, Kif2b and MCAK (Kif2c) (Hirokawa and Tanaka, 2015). There is only one member of the Kif2 family in the ascidian (*P. mammillata*: PmKif2 and *C. intestinalis*: CiKif2) and other non-vertebrate deuterostomes (sFig 2), suggesting that the vertebrate family of proteins evolved from a non-vertebrate deuterostome Kif2a/b/c (henceforth Kif2). Consistent with this ascidian Kif2 also localizes to spindle poles (like Kif2a/Kif2b) and chromosomes (like Kif2c/MCAK) (Figure 3) although CAB localization was strongest. Overexpressing Kif2 to high levels shortened microtubules consistent with it acting as a microtubule depolymerase (sFig 3). In two species of ascidian (*Phallusia mammillata* and *Ciona intestinalis*) Kif2 is a CAB-resident protein (Figure 2 A and sFig 4). By Western blot Kif2 antibody recognizes both endogenous Kif2 and injected Kif2 coupled to GFP (Figure 2 B). Finally, exogenous Kif2::Tomato localized to the CAB during UCD at the 8-cell, 16-cell, and 32-cell stage in *Phallusia* embryos (Figure 2 C).

**Figure 2.**
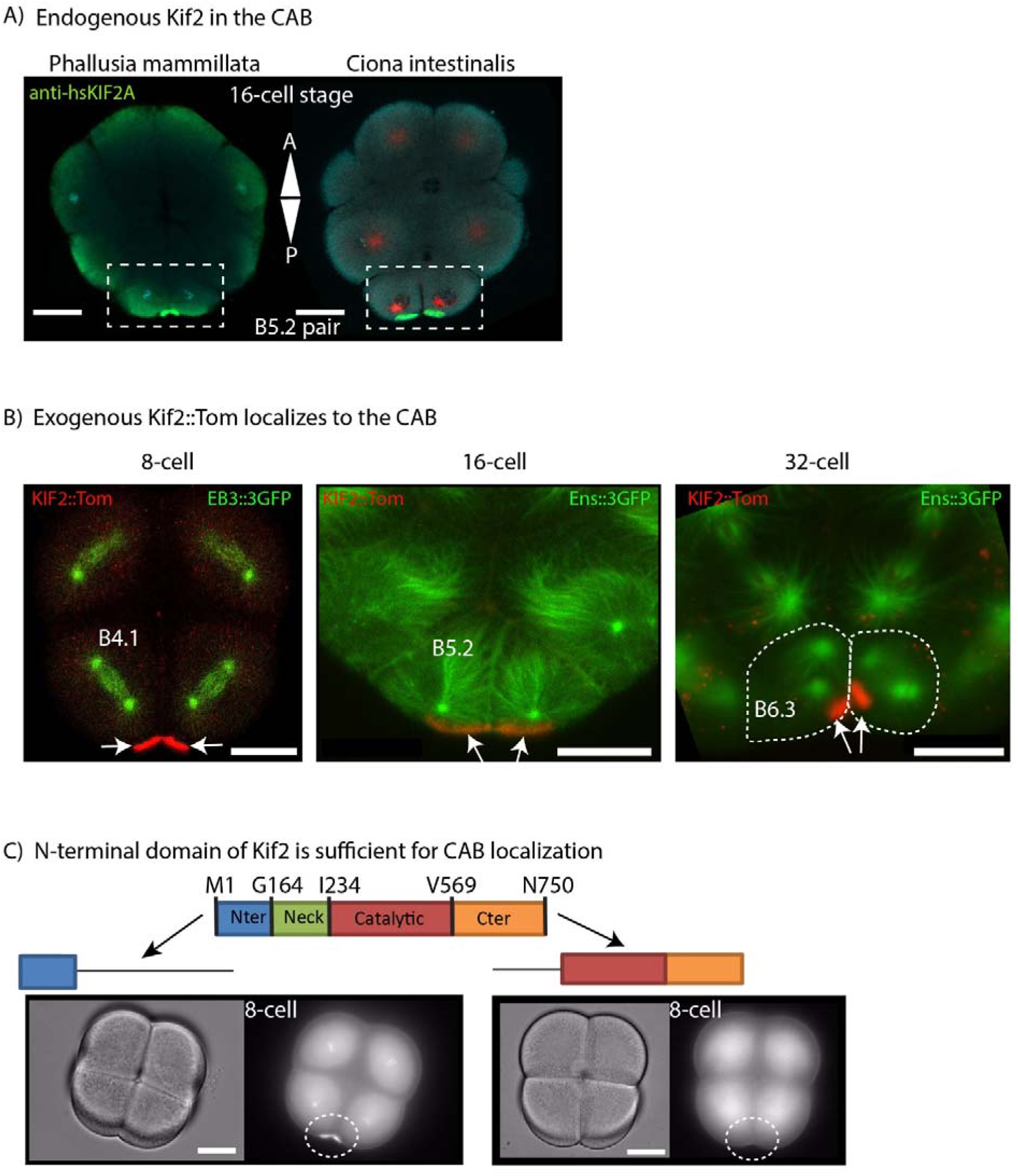
Kif2 protein localizes to the CAB. (A) Endogenous Kif2 in the CAB. Confocal images of two fixed 16-cell stage embryos stained with anti Kif2 from two species of ascidian: *Phallusia mammillata* and *Ciona intestinalis* showing the accumulation of endogenous Kif2 protein (green) in the blastomeres (B5.2 pair) containing the CAB (boxed). DNA stained with DAPI (blue), microtubules with anti-Tubulin (red). n = 94 embryos. Scale bars = 30µm. (B) Western blot. Western blot showing that Kif2 protein is present in unfertilized eggs. Extracts of unfertilized *Phallusia* eggs that were either uninjected (-) or injected (+) with mRNA encoding Kif2::GFP and probed with anti-Kif2 or anti-GFP (left), Ponceau S showing proteins (center) and schematic showing labelling pattern (right). Green bar represents the Kif2::GFP which migrates at a higher molecular mass than endogenous Kif2 protein (pink bar). (C) Exogenous Kif2::Tomato localizes to the CAB. Live *Phallusia* embryos expressing Kif2::Tom (red) and the microtubule markers EB3::GFP (green, 8-cell stage) or Ens::3GFP (green, 16-cell stage and 32-cell stage) showing the localization pattern of Kif2::Tomato to the CAB (red). Arrows indicate CABs. Scale bar = 30µm (8-cell), 30µm (16-cell) and 20µm (32-cell). n = 30. (D) N-terminal domain of Kif2 is sufficient for CAB localization. Schematic of Kif2 protein showing N-terminal domain, neck region, catalytic domain and C-terminal domain. In order to determine which part of the protein was required for CAB localization different truncated versions of Kif2 were fused to Venus and their localization to the CAB followed by epifluorescence. Left: Brightfield and epifluorescence images of 8-cell stage embryo expressing N-ter Kif2::Venus (truncated to amino acids 1–72). Right: Brightfield and epifluorescence images of 8-cell stage embryo expressing truncated N-ter Kif2::Venus (amino acids 239–726). CAB region is indicated by oval. Scale bar = 30µm. n = 16.

**Figure 3.**
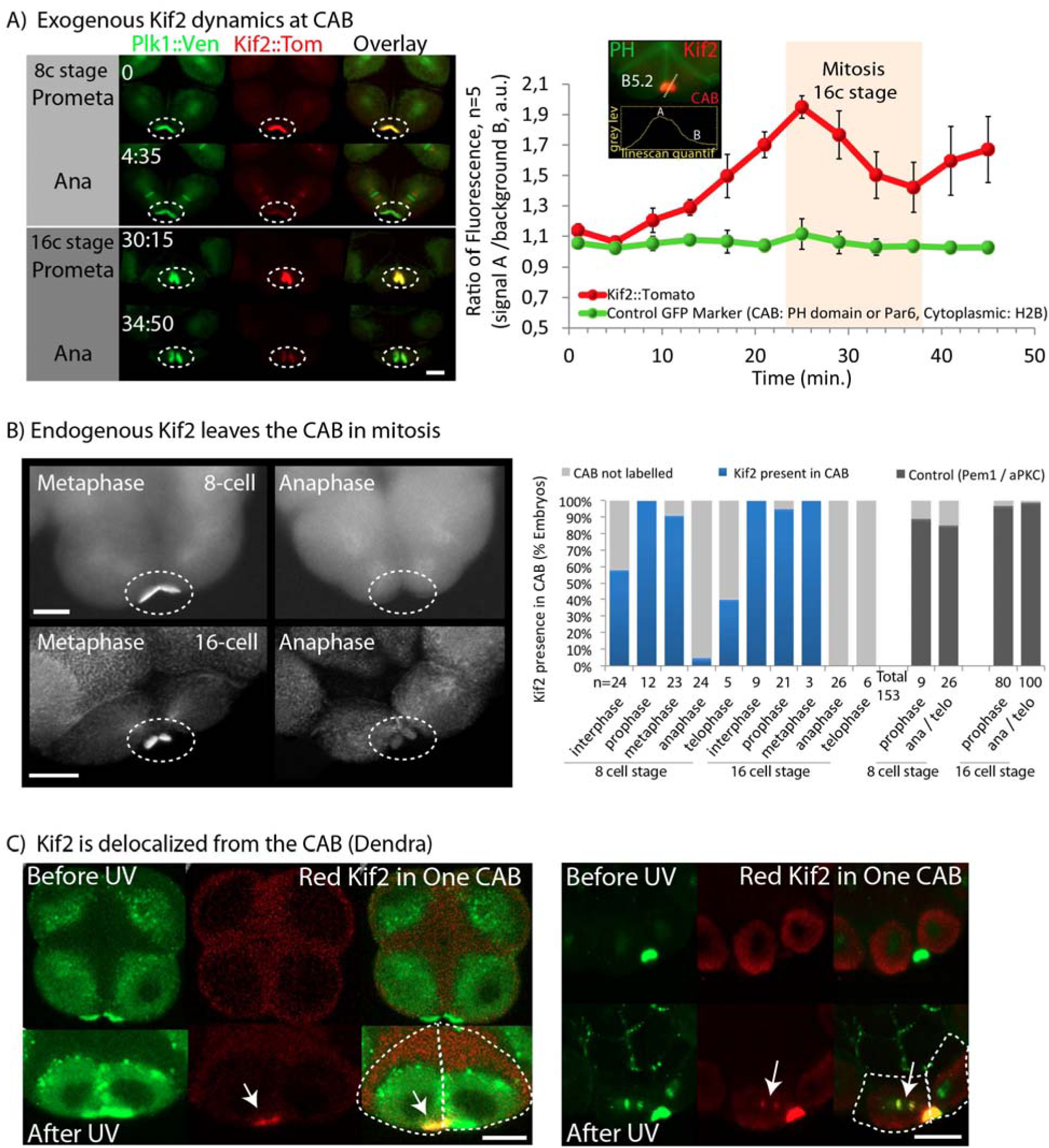
Kif2 dynamics at the CAB. (A) Exogenous Kif2 dynamics at CAB. Unfertilized eggs were microinjected with mRNA encoding Kif2::Tom and Plk1::Venus which also labels the CAB. Confocal images from a time-lapse series showing Plk1::Ven, Kif2::Tom localization at prometaphase and anaphase at the 8 and 16-cell stages. Time in min. Oval indicates CAB region. n = 6. Scale bar = 20µm. Graph to the right shows quantification over one cell cycle of Kif2::Tom fluorescence (reed) versus non-cycling proteins PH::GFP, Par6::Ven or H2B::GFP (green, n=5). Data indicates ratiometric fluorescence signal (linescan : as shown in the inset) in the CAB (mean +/- SEM). (B) Endogenous Kif2 leaves the CAB in mitosis. Fixed 8-cell and 16-cell stage embryos probed with anti-Kif2 during metaphase (left) and anaphase (right); CAB region outlined by oval. Scale bars = 20µm. Quantification of the immunofluorescence data at the 8–16 and 16–32 cell stages (cell cycle stages are indicated). The 4 bars at right are controls showing that two other CAB resident proteins (PEM1 and aPKC) remain in the CAB at Anaphase/Telophase. n is indicated below each column. (C) Kif2 is delocalized from the CAB (Dendra). Embryos at the 8-cell stage containing Kif2::Dendra. Left: Upper row of images show Kif2::Dendra localization at the CAB (Before UV photo-conversion, green). Note absence of red Kif2 at the CAB. Following a brief UV illumination in a pre-defined region of interest centered in only one CAB, the photo-converted red Kif2 is created in one CAB (lower row: After UV, red image and overlay. Arrows indicate CAB containing red Kif2). Right: The second example shows that some red Kif2::Dendra in one CAB diffused away from the CAB to label the nearby chromosomes (arrows). The chromosomes in the adjacent blastomere are not labelled with the red Kif2::Dendra. Scale bars = 30µm. n = 6

Next we wished to pinpoint which domain of Kif2 protein was required for CAB localization. For this we injected mRNAs encoding fusions between Venus tag and various portions of Kif2 and evaluated CAB localization of each construct. Our results show that the N-terminal 72 amino acids of Kif2 was capable of driving localization to the CAB, and conversely removing the N-terminal domain abolished CAB localization of the Venus-tagged constructs (Figure 2 D).

### Kif2 cycles onto and off of the CAB

We noticed that Kif2 protein appeared to accumulate on the CAB during interphase and leave the CAB during mitosis. In order to confirm that Kif2 cycled onto and off of the CAB we performed live cell ratiometric imaging of Kif2::Tom levels relative to Plk1::Ve or other fluorescent constructs (Par6::Venus, H2B::GFP or PH::GFP). Our analysis revealed that Kif2 protein accumulates at the CAB during interphase peaking at early mitosis and that Kif2 protein begins to leave the CAB during prometaphase with levels reaching a minimum at late anaphase/cytokinesis onset (Figure 3 A and Movie S7). Careful examination of endogenous Kif2 levels at the CAB in either *Phallusia* or *Ciona* embryos revealed a similar dynamic delocalization of Kif2 during mitosis, whereby endogenous KIF2 protein was delocalized from the CAB by late anaphase (Figure 3 B and sFig 4). Other CAB-resident proteins such as PEM1 or aPKC do not leave the CAB during mitosis (Figure 3 B). In order to determine whether loss of signal at the CAB was due to delocalization or degradation we used the photo-convertible construct Kif2::Dendra to follow a specific pool of Kif2 protein *in vivo*. UV illumination converts the green Kif2::Dendra fusion protein into a red one. UV illumination of Kif2::Dendra in a region within one CAB caused just one CAB to become red (Figure 3 C). Since Kif2 is also a kinetochore localized protein, we reasoned if we could detect red Kif2::Dendra on chromosomes that would indicate that the red version of Kif2 protein left the CAB following photo conversion to become captured by adjacent chromosomes (it is important to note that this does not rule-out destruction but does show that red Kif2 protein leaves the CAB). Figure 3 C shows that we were able to detect red Kif2::Dendra on chromosomes in the blastomere containing the photo-converted red Kif2::Dendra. Thus the red Kif2 protein we detected on the chromosomes came from the CAB (Figure 3 C), indicating that some Kif2 protein leaves the CAB in mitosis.

### Kif2 localizes to cortical ER in the CAB

As noted previously, the CAB is a multilayered structure, comprised of a thick layer of cortical endoplasmic reticulum (cER) protruding into the cytoplasm which adheres to a specialized region of actin-rich cortex (Patalano et al., 2006). By co-staining for mitochondria which surround and outline the cortical ER mass, and for pMNK, a cER resident protein (Paix et al., 2011b), the surface and deeper cER can be visualized (Figure 4 B). Unlike aPKC protein which is contiguous with the actin layer, Kif2 protein occupies the thicker cER layer (Figure 2). To determine the precise localization of Kif2 protein in the CAB we prepared isolated cortices. By sticking 8-cell stage embryos to coverslips followed by shearing using an isotonic buffer, the cortex and its associated cER is retained on the cover slip (Figure 4 A). Probing these cortical preparations with anti-Kif2 revealed a concentration of Kif2 in the cER of the CAB (Figure 4 A, and inset for higher definition). In live embryos we were able to observe dynamic finger-like projections from the surface of the CAB labelled with Kif2::Tom (Figure 4 C), consistent with the notion that Kif2 was localized to the tubes of cER in the CAB.

**Figure 4.**
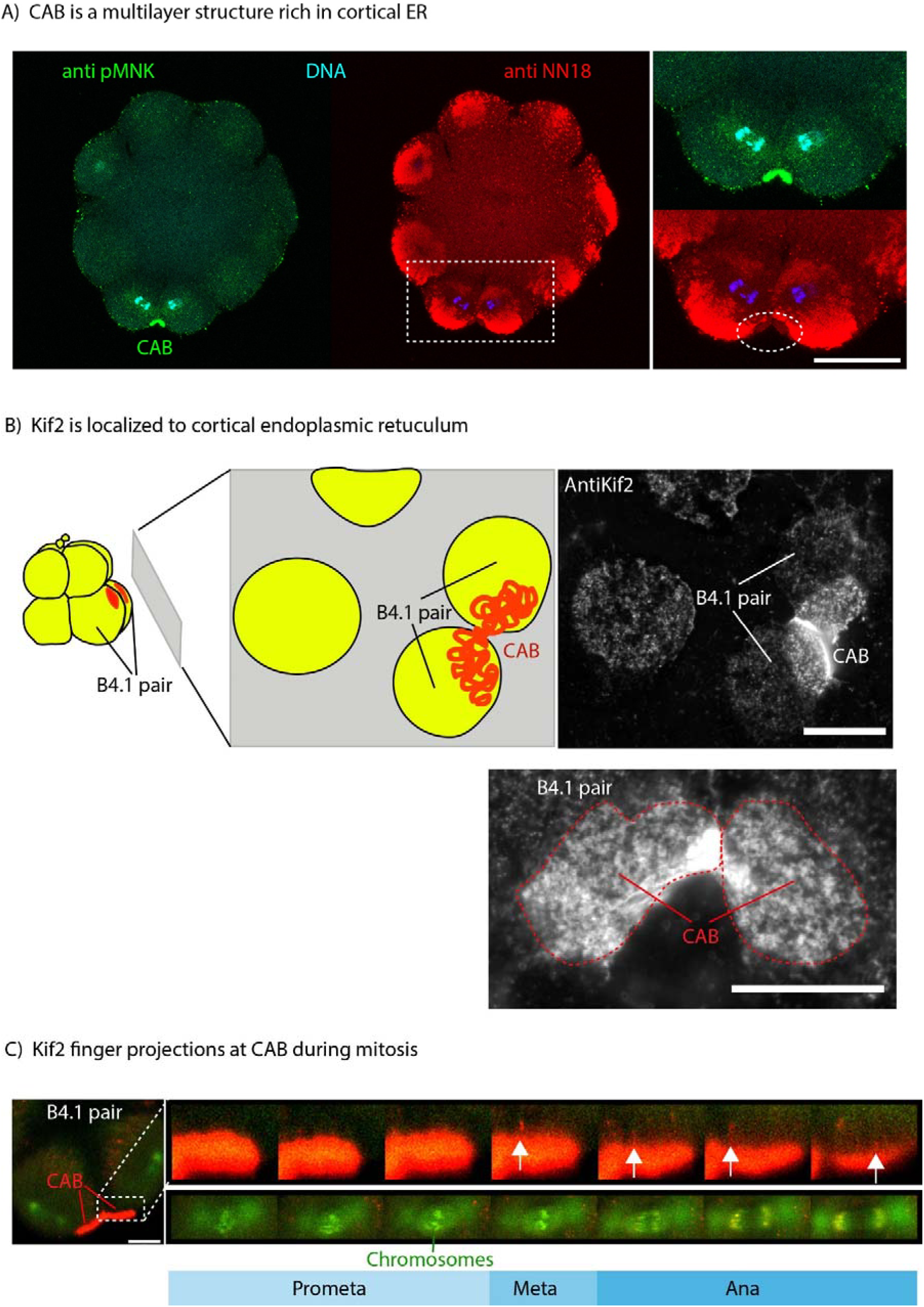
Kif2 protein localizes to the domain of cortical ER in the CAB. (A) CAB is a multilayered cortical structure rich in cortical ER. pMNK labelling of the cortical ER (green) with the mitochondria labelled with anti NN18 (red). Enlarged view of the ROI showing that the mitochondria are excluded from the CAB where the cER is enriched (oval ROI). Scale bar = 30µm. (B) Kif2 is localized to cortical endoplasmic reticulum. Schematic of cortical preparation, ER in red. Probing cortical preparations with Kif2 antibody revealed that Kif2 protein was localized to the domain of cortical ER in the CAB. Enlarged view of an 8-cell stage cortical preparation showing the cER labelled with anti-Kif2. CAB is indicated in boxed region. Scale bars = 20µm. (C) Kif2 finger-like projections at CAB surface during mitosis. Embryo containing Kif2::Tom, EB3::GFP and H2B::GFP. During mitosis red finger-like projections appear at the surface of the CAB (arrows). Mitotic stage is indicated from H2B::GFP labelling of chromosomes and Kif2::Tom to kinetochores during mitosis. The inset shows the ROI together with the cell cycle stage as defined by the chromosome behavior. Selected images from a timelapse. Scale bar = 20 µm.

### Microtubule dynamics at the CAB

To visualize the rapid dynamics of microtubule at the cortex embryos have to be immobilized close to a flat surface, here provided by the coverslip. In one cell *C.elegans* zygotes this permitted the accurate determination of plus end dynamics leading to the development of the “touch and pull” mechanism of ACD (Kozlowski et al., 2007). In ascidians, measurement of microtubule plus end dynamics in a cortical slice containing the CAB and non-CAB cortex is complicated by both the movement and the geometry of the embryo: since the CAB is an apical cortical structure close to the midline it invariably curves away from the coverslip. In order to overcome these problems we dissociated blastomeres with calcium-free seawater and used protamine-coated coverslips to immobilize them (Figure 5 A). Blastomere isolation does not perturb UCD in *Phallusia* embryos (Prodon et al., 2010). Ens::3GFP fluorescence was then measured in a cortical slice within 1µm of the coverslip every 0.8 seconds (Figure 5).

**Figure 5.**
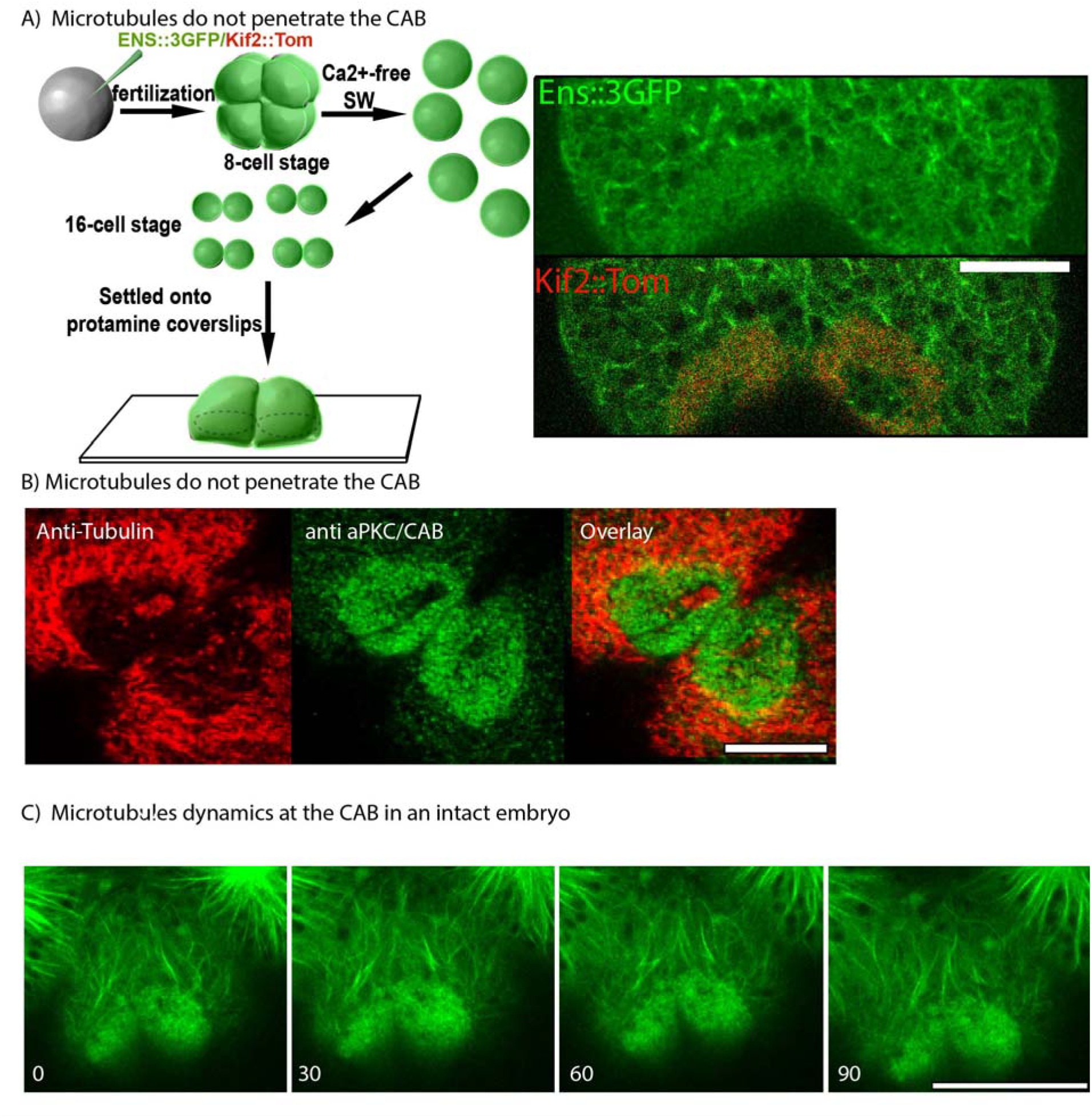
Microtubule dynamics at the cortical CAB. (A) Microtubules do not penetrate the CAB. In order to follow microtubule dynamics at the CAB unfertilized eggs were microinjected with mRNA for Ens::3GFP and Kif2::Tom, fertilized and then transferred to calcium-free seawater at the 8-cell stage to dissociate the 8 blastomeres. Once the dissociated blastomeres had divided, pairs of B5.2 blastomeres were placed on coverslips that had been treated with protamine so that the blastomeres adhered. Next fast confocal imaging (0.8 sec/image) of a 1µm thick optical section just above the coverslip revealed that microtubules were absent from the CAB (red). Scale bar = 10 µm. n = 4. See Movie S7. (B) Microtubules do not penetrate the CAB. Confocal z-sections from fixed 8-cell stage embryos showing microtubules (red, anti-tubulin) and the CAB (green, anti-aPKC). Microtubules are absent from the CAB (n>50). Scale bar = 20µm. (C) Microtubule dynamics at the CAB in an intact embryo. Selected frames from a 32-cell stage embryo previously injected with mRNA encoding Ens::3GFP and Kif2::Venus. Confocal stacks were acquired every 30 seconds and selected frames from each stack are shown. The CAB is visible since it accumulates Kif2::Venus. Microtubules do not penetrate the CAB during all of interphase (see Movie S8). Scale bar = 20µm.

Microtubules are present as a dense network on the cortex during interphase but in the CAB domain they are absent (Figure 5 A, B, see Movie S8). In Movie S8, microtubules can be observed polymerizing in the direction of the CAB (labelled red with Kif2::Tom) but they fail to penetrate the CAB. Occasionally during interphase the CAB has a CAB-free zone at its center and microtubules can be seen to reach the cortex in the hole in the center of the CAB but are absent from the Kif2-labelled CAB (Figure 5 A, B). Microtubules also avoid the CAB in the intact embryo at the 32-cell stage even though they can be seen growing around the CAB (Figure 5 C and Movie S9) as in isolated blastomeres (Figure 5 A), however since the embryo moves in the imaging plane many z sections were acquired so the time resolution in one z plane is greatly reduced.

### Microtubule depolymerization and aster asymmetry

In order to determine the role of Kif2 in the CAB we generated a mutant form of Kif2 protein (DN-Kif2) based on the construct which behaves as a dominant negative in mammalian cells (Wordeman et al., 2007). DN-Kif2 contains the N-terminal domain which targets it to the CAB and the C-terminal domain which permits dimerization, but lacks the catalytic domain (Figure 6). mRNA encoding the DN Kif2 and H2B::Rfp1 constructs were injected into one blastomere of a two cell stage embryo previously injected with Ens::3GFP and Kif2-Nter::Venus to monitor microtubules and the CAB in both halves of the embryo (Figure 6). DN Kif2 prevented the development of aster asymmetry and abolished UCD (Figure 6Ai). We measured the distance between the nearest spindle pole and the CAB in both halves of the embryo at the 8-cell stage and found that DN Kif2 significantly increased the spindle pole to CAB distance (Figure 6 Ai). Since taxol stabilizes microtubules and reduces depolymerization we reasoned that taxol should have a similar effect to DN Kif2. Treating embryos in mitosis with taxol also prevented the development of aster asymmetry (Figure 6 Aii). These data suggest that aster asymmetry and asymmetric spindle positioning require the activity of Kif2 and microtubule depolymerization.

**Figure 6.**
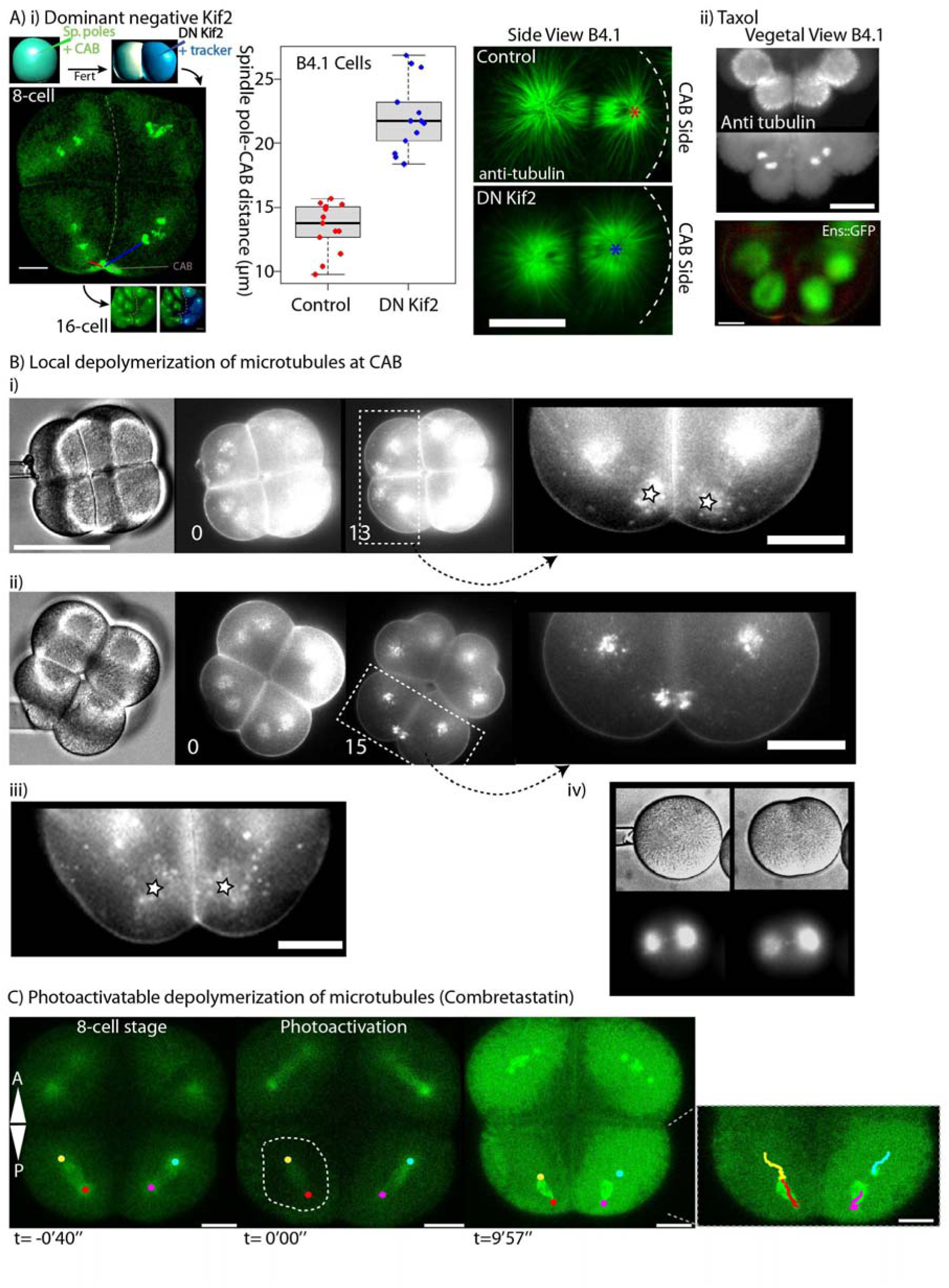
Inhibition of Kif2 and localized microtubule depolymerization. (A) Dominant negative Kif2. (i) Left: 8-cell stage embryo in which one blastomere at the 2-cell stage was microinjected with a truncated version of Kif2 which acts as a dominant negative (henceforth DN-Kif2) and histone H2B::Rfp1 as a fluorescent tracker to label the injected half of the embryos. Unfertilized eggs had previously been injected with Ens::3GFP to monitor spindle poles and Kif2 Nter::Ven to monitor the CAB. Center: Minimum spindle pole distance to the CAB was measured at cleavage onset in the control half of the embryo (blue line) and the half containing DN-Kif2 (red line). Right: Embryos injected at the 2-cell stage were also fixed and labelled for microtubules. A side view of an 8-cell stage embryo shows that the CAB-proximal aster is larger in the presence of DN-Kif2, and the spindle pole distance to the CAB cortex is increased (stars represents the center of the spindle pole closest to the CAB). (ii) Treating embryos at the 8-cell stage with taxol to stabilize microtubules also created equal sized asters. Top: Epifluorescence images of fixed 8-cell stage embryo showing microtubules (anti-Tubulin) and DNA (DAPI). Lower: Epifluorescence image of live 8-cell stage embryo treated with taxol showing merged images (microtubules green, CAB red). Scale bars = 20µm. (B) Local depolymerization of microtubules at CAB. (i) Nocodazole pipette (small bore) was advanced towards one B4.1 blastomere (bright-field image time 0). Spindle position during mitosis (13 min. image). Zoom of boxed region showing the spindle pole closest to the pipette moved even closer towards the CAB during mitosis (compare left spindle with right spindle pole – stars). See Movie S10. (ii) Nocodazole pipette (large bore) was advanced towards one B4.1 blastomere (bright-field image time 0). Fluorescence image showing spindle position during mitosis (15 min. image). Zoom of boxed region showing that both spindle poles moved closer to the CAB and midline (arrows). All scale bars are 30 µm. (iii) Control blastomeres at 8-cell stage. (iv) Nocodazole pipettes were tested on fertilized eggs containing Ens::3GFP. Note loss of microtubule density in the aster nearest the pipette. Time in min. All scale bars = 30µm. See Movie S11. (C) Microtubule depolymerization with the caged Combretastatin. Caged Combretastatin was uncaged causing its activation in the boxed region causing microtubule depolymerization during mitosis. Microtubules were labelled with Ens::3GFP. Spindles were left intact and still migrated towards the CAB. Due to diffusion of the uncaged Combretastatin all cells were affected. Tracking of spindle poles is shown in the inset to the right. Scale bars = 20µm.

If aster reduction facilitates the migration of one spindle pole towards the cortex, we reasoned that increasing microtubule depolymerization near the CAB would enhance spindle pole movement towards the CAB, since microtubules would become shorter. We employed a method to depolymerize microtubules locally by using a micro-pipette as a spatially-confined source of nocodazole (see materials and methods) (Figure 6 Bi and Movie S10 and S11). Since it was not possible to perform these experiments on our confocal microscope we used an epifluorescence system. We bathed embryos in Cell Mask orange which labels both the plasma membrane and vesicles that accumulate around spindle poles during mitosis (McDougall et al., 2015). This was more convenient that using Ens::3GFP and also gave better visualization of both spindle poles in each blastomere. However, to confirm that these nocodazole micropipettes caused localized microtubule depolymerization we imaged microtubules in zygotes (Figure 6 B iv). We applied different diameter pipettes with the same nocodazole concentration near the CAB in one blastomere at the 8-cell stage and measured the effect on the position of the proximal spindle pole. Relatively small diameter nocodazole pipettes caused the proximal spindle pole to migrate closer toward the CAB and midline than normally observed (Figure 6 Bi and Movie S10). Importantly, the blastomeres far from the source of nocodazole were unaffected and divided normally (Figure 6 Bi and Movie S10). By using a larger bore pipette we could affect both CAB-containing blastomeres so that both CAB-proximal spindle poles moved closer to the CAB and midline while blastomeres farther from the pipette behaved normally and divided (Figure 6 Bii). Control DMSO-containing pipettes had absolutely no effect on spindle positioning (Figure 6 Biii). To depolymerize microtubules via a second method we used photo-activation of caged Combretastatin while imaging microtubules with Ens::3GFP (Figure 6 C). Activation of caged Combretastatin during mitosis caused astral microtubule depolymerization while leaving spindle microtubules intact. As noted with nocodazole pipettes, the whole spindle migrated even closer towards the CAB (Figure 6 C). These data revealed that reducing aster size could augment the asymmetric positioning of the spindle, supporting the hypothesis that astral microtubule length sets the distance of the spindle pole to the CAB.

## DISCUSSION

Here we present evidence for the localization and function of a microtubule depolymerase (Kif2) at a cortical site that causes UCD. Precise control of microtubule dynamics at the cortex is a fundamental cellular mechanism that is involved in diverse biological processes ranging from the control of axon morphology (Homma et al., 2003; Maor-Nof et al., 2013; Noda et al., 2012) to the unequal division of cells (Cowan and Hyman, 2004; Siller and Doe, 2009). UCD is an extremely widespread process in biology occurring in bacteria (Adams and Errington, 2009; Aldridge et al., 2012), yeast (Carminati and Stearns, 1997), and many different types of embryo. Amongst embryos, UCD has been observed in ctenophores (Martindale and Henry, 1999), chaetognaths (Carré et al., 2002), spiralians (Lambert, 2010), echinoderms (Schroeder, 1987; Holy and Schatten, 1991), and invertebrate chordate embryos of the ascidian (Hibino et al., 1998). A common theme among several embryos that display UCD is the development of aster asymmetry. Unequally cleaving spiralian embryos (*Tubifex*, and the leech *Helobdella*) display aster asymmetry at the 1 to 2 -cell stage leading to eccentric positioning of the mitotic spindle (Weisblat, 2007). Sea urchin embryos also display UCD starting at the 8 to 16-cell stage culminating in the formation of four micromeres (Schroeder, 1987; Holy and Schatten, 1991) at the vegetal pole where aPKC is absent (Prulière et al., 2011). Again the mechanism underlying this UCD is not known in these embryos, however, observations indicate that one centrosome becomes disc-shaped and closely apposed to the cortex before UCD (Schroeder, 1987; Holy and Schatten, 1991). In ascidians three successive rounds of UCD occur starting at the 8 to 16-cell stage, resulting in the formation of two small blastomeres at the 64-cell stage that are germ cell precursors (Hibino et al., 1998). We showed previously that the CAB causes one pole of the mitotic spindle to approach the CAB during prometaphase through anaphase which is accompanied by the shrinking of the CAB-proximal aster (Prodon et al., 2010).

One central unresolved question therefore is how specialized cortical sites affect astral microtubules leading to the development of aster asymmetry. Here we show that in ascidian embryos the microtubule depolymerase Kif2 plays a key role in promoting aster asymmetry which in turn is required for UCD. Kif2 is concentrated on a subdomain of cortical endoplasmic reticulum (cER) found at the centrosome-attracting body or CAB during interphase and leaves the CAB during mitosis (Figure 4). We propose that Kif2 affects local microtubule dynamics at the cortex in the vicinity of its location, causing depolymerization of the nearest microtubules during mitosis thus leading to the development of aster asymmetry. By artificially increasing the amount of local depolymerization of astral microtubules in the proximity of the CAB we found that the spindle pole moved even closer to the CAB (Figure 6).

Our results have led us to the conclusion that polymerization of astral microtubules opposes the pulling forces that likely displace the mitotic spindle towards the CAB. We therefore propose that the diffusion of Kif2 from the CAB in mitosis causes the local depolymerization of those astral microtubules nearest the CAB thus facilitating the eccentric positioning of the mitotic spindle near the CAB cortex.

The mechanism we have discovered here in a chordate deuterostome embryo extends our understanding of UCD, which so far has been heavily studied in two protostomes, *C.elegans* and *Drosophila*. In *Drosophila* larval neuroblasts, even though the apically localized Gα/Pins/NuMA complex aligns the mitotic spindle (Nipper et al., 2007; Siller et al., 2006; Siller and Doe, 2009), UCD can be driven by a cortical subdomain of Myosin II via a mechanism which is independent of spindle position (Cabernard et al., 2010). Work in *C. elegans* zygotes has demonstrated the fundamental importance of the conserved Gαi/Pins/NuMA complex, which accumulates at the posterior cortex and creates a pulling force on astral microtubules that is greater at the posterior cortex, thus causing unequal cleavage (Grill et al., 2001; Redemann et al., 2010). Intriguingly, microtubule depolymerization is thought to be required for UCD in *C. elegans* (Gönczy et al., 2001; Kozlowski et al., 2007; Nguyen-Ngoc et al., 2007) although the mechanism responsible is currently unknown. It appears that this is not Kif2-dependent, since inhibition of MCAK with RNAi does not prevent eccentric spindle positioning (Grill et al., 2001).

The control of microtubule plus end dynamics during UCD by Kif2 is remarkably similar to the axonal pruning function of Kif2 in mammalian post-mitotic neurons. For example, in developing neurites Kif2A is thought to bind to the plasma membrane associated protein phosphatidylinositol 4-phosphate 5-kinase (Noda et al., 2012). Kif2A localizes to the tips of developing neurites *in vivo* and functions to suppress collateral branch extension since the knockout of Kif2A in mice causes increased microtubule stability at the cell edge in growth cones creating more growth of collaterals (Homma et al., 2003).

Our results presented here indicate that the flattened aster with short microtubules and movement of the spindle toward the CAB are both mediated by the action of a cortically-localized microtubule depolymerase. Given that Kif2 controls UCD in invertebrate chordate embryos and morphogenesis of developing mammalian neurites, it may be worthwhile investigating the role played by kinesin-13 family members such as Kif2 and other microtubule depolymerases during UCD as well as other biological processes where microtubule dynamics alter when they encounter the cortex.

## METHODS

### Origin of the animals

*Phallusia mammillata* were collected at Sète (Etang de Tau, Mediterranean coast, France) and *Ciona intestinalis* at Roscoff. Ascidian gamete collection, dechorionation, fertilization and embryo cultures were as described previously (McDougall et al., 2014).

### Antibodies, fixation and reagents

Embryos were fixed in −20° methanol containing 5uM EGTA and immunolabelled as previously described (Sardet et al 2011). We used anti aPKC (Santa Cruz 216) and anti PEM1 (Paix et al., 2011b) to label the CAB, anti-tubulin (YL1/2 and DM1a, Sigma) for microtubules, anti Kif2 (Ganem and Compton, 2004), γ-tubulin (Sigma GTU88) for centrosomes. DiI and DiO (Invitrogen) were used to label ER and mitochondria respectively, Paclitaxel (Sigma) to stabilize microtubules and nocodazole (Sigma), caged Combretastatin (provided by D. Burgess) to depolymerize microtubules.

For Western blot samples were prepared in Laemmli sample buffer and migrated on 10 % polyacrylamide gels by SDS-PAGE using standard procedures. Two lanes were loaded containing either 40 uninjected eggs or 40 eggs which had been injected with mRNA encoding full-length Kif2::GFP. After transfer to nitrocellulose, each lane was cut into two strips lengthwise which were incubated overnight with either anti-Kif2 or anti-GFP at a dilution of 1:1000 in TBS + 5% dry milk. The 4 strip blots were then washed in TBS + 0.1% tween, incubated with anti-rabbit secondary at 1:10000, and washed in TBS-tween. The signal was detected using West Pico chemiluminescent substrate (Fisher) at an exposure of 2 minutes.

### Preparation of isolated cortices

Isolated cortices were prepared following the methods previously established in the laboratory (Patalano et al., 2006; Sardet et al., 2011). Briefly, embryos at the desired stage are transferred to calcium free seawater then placed at high density on cover slips coated with protamine (1 mg/ml) to which they adhere within 1 minute. After two washes with cortex isolation medium (CIM (cortex isolation medium): 0.8 M glucose, 0.1 M KC1, 2 mM MgCl2, 5mM EGTA, 10 mM MOPS buffer, pH 7), a stream of CIM is sprayed gently with a Pasteur pipette, shearing off the embryos but leaving attached to the glass imprints of the adherent membrane and associated cortical structures. The coverslips are washed rapidly in CIM then placed in cold methanol for fixation, then rehydrated in PBS and processed for immunofluorescence by standard procedures used for whole embryos.

### Microinjection and imaging

Microinjection was performed as previously described (McDougall et al., 2014). Briefly, dechorionated oocytes were mounted in glass wedges and injected with mRNA (1–2 µg/µl pipette concentration/ ~2% injection volume) using a high pressure system (Narishige IM300). mRNA-injected oocytes were left for 2–5 hours or overnight before fertilization and imaging of fluorescent fusion protein constructs. The lipophilic dye Cell Mask Orange (Molecular Probes) was prepared at a concentration of 10 mg/ml in DMSO and diluted in sea water at 20 µg/ml then mixed 1:1 with the embryos just prior to imaging. Epifluorescence imaging was performed with an Olympus IX70, Zeiss Axiovert 100 or Axiovert 200 equipped with cooled CCD cameras and controlled with MetaMorph software package as previously described. Confocal microscopy was performed using a Leica SP5 or SP8 fitted with 40x/1.3na oil objective lens and 40x/1.1na water objective lens. All live imaging experiments were performed at 18–19°C. For fast imaging of cortical preparations a rectangular image section of the imaging array was selected to increase the temporal resolution to 0.8 images/sec. Image analysis was performed using Image J, ICY and MetaMorph software packages. Calcium-free sea water: 450 mM NaCl, 9 mM KCl, 33 mM Na2SO4, 2.15 mM NaHCO3, 10 mM Tris pH 8, 2.5 mM EGTA.

### Micromanipulation

All manual micromanipulation experiments were performed on an Olympus IX70 microscope using a 20x objective lens and Metamorph acquisition software. Embryos at the 4-cell stage were incubated with Cell Mask Orange diluted in sea water (1/1000) for 90 sec. then washed with sea water. Cell Mask labelled embryos were mounted at the 8-cell stage for observation. To prepare the nocodazole pipettes, nocodazole was added to liquid 1% low melt agarose in sea water giving a final concentration of 50µM nocodazole. Using a Narishige PN30 puller microinjection pipettes were pulled from GC100-T glass (filament-free) capillary tubes. The tips of the microinjection pipettes were broken and calibrated by microscopic observation. These micropipettes were dipped into the liquid nocodazole/agarose solution, placed at room temperature which caused the agarose containing nocodazole to solidify. These prepared microinjection needles were stored in humid chambers and used the same day. The nocodazole/agarose needles were advanced towards the 8-cell stage embryos and placed on the surface of one B4.1 blastomere near the CAB starting at nuclear envelope breakdown. Bright-field and fluorescence images were acquired every 10 seconds using MetaMorph software package.

### Synthesis of RNAs

We used the Gateway system (Invitrogen) to prepare N and C terminal fusion constructs using pSPE3::Venus (a gift from P. Lemaire), pSPE3::Rfp1, pSPE3::Cherry, pSPE3::tomato for all constructs except PH::GFP which was cloned into pRN3. For construct details please refer to our previous methods publication (McDougall et al., 2015) All synthetic mRNAs were transcribed and capped with mMessage mMachine kit (Ambion).

### Bioinformatics

We created a database of animal protein sequences derived from the complete genomes of various metazoan lineages: *Amphimedon queenslandica*, *Mnemiopsis leidyi*, *Trichoplax adhaerens, Nematostella vectensis, Acropora digitifera, Hydra vulgaris, Crassostrea gigas, Aplysia californica, Capitella teleta, Lingula anatina, Caenorhabditis elegans, Drosophila melanogaster, Strongylocentrotus purpuratus, Branchiostoma floridae, Ciona intestinalis, Phallusia mammillata* and *Homo sapiens*. All data was retrieved from the NCBI genomes portal: https://www.ncbi.nlm.nih.gov/genome/browse/

We searched this database with the PFAM Kinesin hidden Markov model [PMID: 26673716] using the global local strategy implemented in HMMER2 [http://hmmer.org/download.html] and the model specific 'gathering threshold' bit scores as a cutoff. Kinesin regions (649 sequences), as defined by HMMER2 alignments, were extracted from full length sequences and aligned using the MAFFT software package with default parameters [PMID: 12136088].

This alignment was used to create a phylogeny of all Kinesin domains, using the Bayesian approach implemented in phylobayes, with an LG+G model of sequence evolution [PMID: 19535536]. Two chains were run for 3500 generations. 700 generations were discarded as burnin. Although the chains had not converged, the region of the phylogeny around the human Kif2 proteins revealed a stable clade composed of orthologs of the human KIF19, KIF18, KIF24 and KIF2A/B/C genes with full support from posterior probabilities. The sequences representing this clade of orthologous groups were extracted. The *Phallusia mammillata* Kif13 protein sequence was added to the alignment, and another phylogeny reconstructed using the Phyml package with an LG+G evolutionary model and 100 bootstrap replicates [PMID: 20525638]. This tree is shown in sFig. 2.

## ACKNOWLEDGEMENTS

We thank Duane Compton and Bernardo Orr for the generous gift of anti Kif2 antibody, Linda Wordermann for useful advice, Remi Dumollard for thoughtful insights, and members of the LBDV for helpful discussions. We thank Mafalda Loreti and Paul Stolz for preliminary experiments that do not appear in the article. This work was supported by the Agence National de la Recherche (ANR-12-BSV2-0005-01), and in part by support from Sorbonne Universities ANR-11-IDEX-0004-02 to the Picard Network, and by NSF grant 124425 to DB.

## AUTHOR CONTRIBUTIONS

A.McD, J.C. and V.C conceived project and conducted experiments.

A.McD and J.C. wrote the ms.

V.C, G.P, D.B, and M.F performed experiments.

R.C performed bioinformatics analysis.

C.H and L.B provided molecular tools.

## COMPETING FINANCIAL INTERESTS

The authors declare no competing financial interests.

---

**Figure S1.**
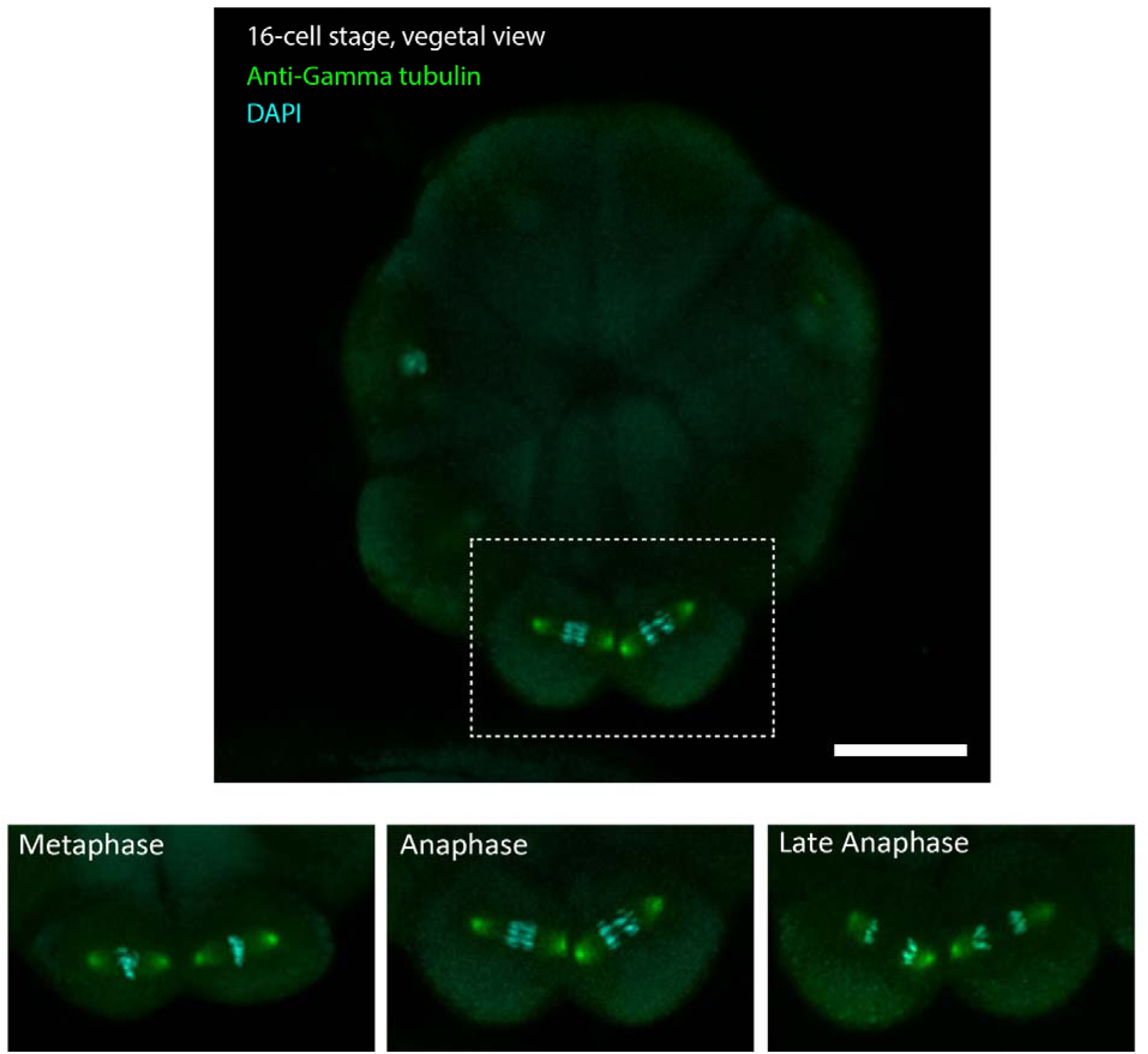
Gamma-tubulin presence at spindle poles during mitosis. 16-cell stage *Phallusia mammillata* embryos fixed and labelled with anti-γ-tubulin (green) antibody and DAPI (blue). Inset showing γ-tubulin at metaphase, anaphase and late anaphase of the posterior pair of blastomeres (B5.2). γ-tubulin is present at both spindle poles throughout mitosis. Scale bar = 30µm.

**Figure S2.**
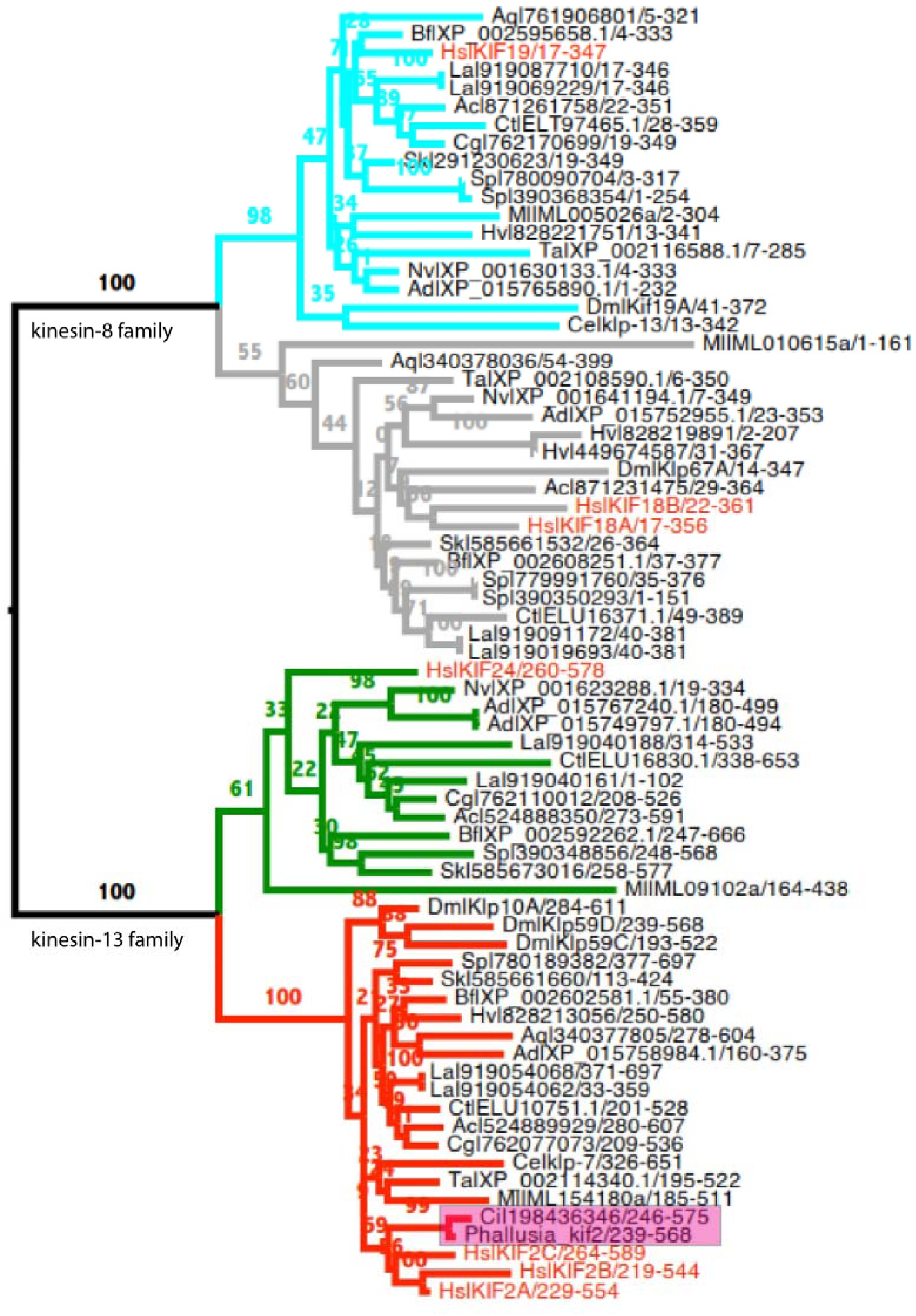
Phylogenetic analysis of Kif2 (kinesin-13) family of proteins. Phyml tree of the clade of animal KIF19, KIF18, KIF24 and KIF2A/B/C ortholog groups. Kif2 and Kif24 (bottom half of tree) are members of the Kinesin-13 family of microtubule depolymerases; Kif18 and Kif19 (top half of tree) are members of the related Kinesin-8 family (Lawrence et al., 2004). With the exception of the *Phallusia* sequence, the first two letters of leaf names represent the species of the sequence (see methods for list). These are followed by identifiers retrievable from the NCBI protein sequence database. Bootstrap support is show above branches. Although within the gene groups the species phylogeny is generally not well resolved, there is high support for lineage specific duplications of Kif2 in *Drosophila* and Humans, with the majority of invertebrate taxa having only a single representative, strongly suggesting the state in the ancestral animal was a single gene. *Ciona* and *Phallusia* data are highlighted by the pink box.

**Figure S3.**
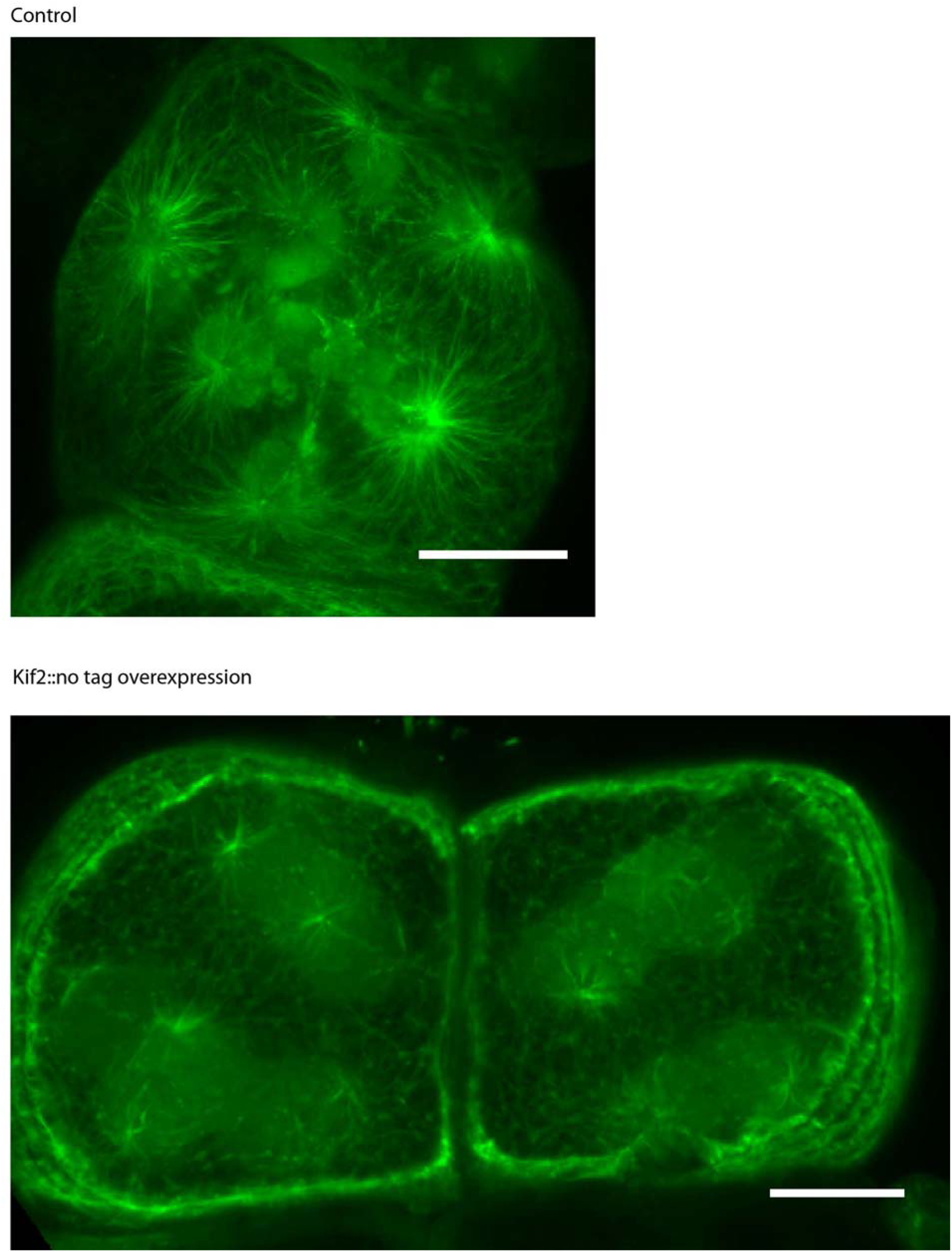
Microtubule density is reduced in cells overexpressing Kif2. Eggs injected with mRNAs encoding Kif2 (without a fluorescent tag) and Ens::3GFP or only Ens::3GFP (top) were fertilized then treated with cytocholasin B to prevent cell division. Microtubule density was greatly reduced in the presence of excess Kif2. Scale bar = 20µm.

**Figure S4.**
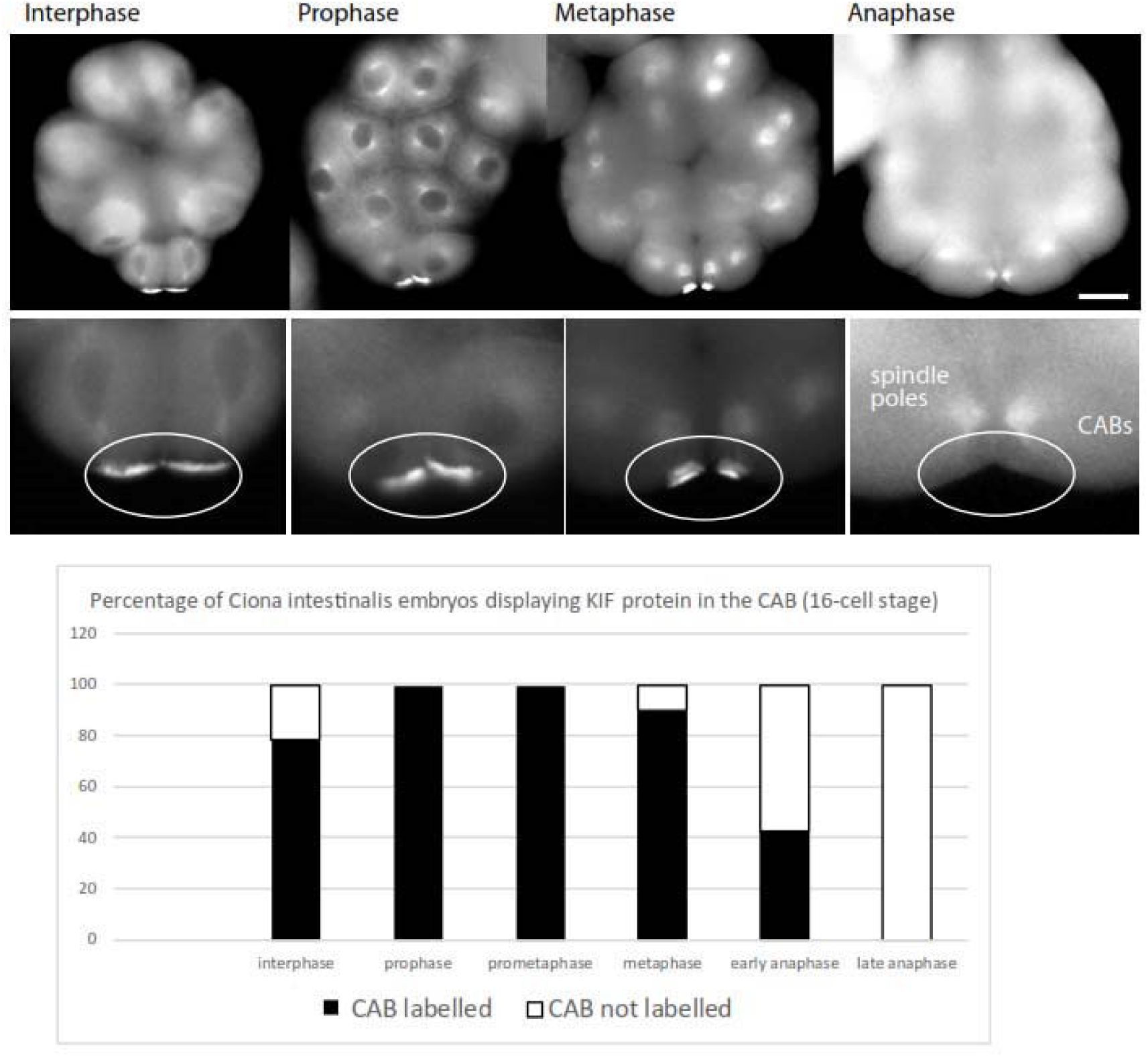
Kif2 protein is delocalized from the CAB in Ciona intestinalis during mitosis. Immunolabelling with anti-Kif2 of 16-cell stage *Ciona* embryos. Kif2 labels the CAB during later interphase/early prophase and is absent at the CAB during anaphase. Scale bar = 30 µm. Graph showing percentage of embryos (total n=94) with CAB labelled at different stages of the cell cycle.

## Supplementary Movies (please go to: http://movincell.org/)

### Movie S1. Unequal cell division from 8 to 16-cell stage

Bright-field images of 8-cell stage *Phallusia mammillata* embryo viewed from the vegetal side from interphase until cell division. The two vegetal posterior blastomeres that divide unequally (B4.1 pair) are at the bottom. Following nuclear envelope breakdown it is possible to discern the movement of the mitotic spindle in the two bottom blastomeres. Time in min:sec.

### Movie S2. Unequal Cell Division 1. 8-16-cell

Unfertilized *Phallusia* eggs injected with Ens::3GFP (green). Confocal images of 8-cell stage embryos showing microtubule behavior during late interphase and mitosis. B4.1 pair of blastomeres are at the bottom. Note that the aster is asymmetric in both bottom blastomeres. Time in min. Scale bar = 30µm.

### Movie S3. Unequal Cell Division 2. 16 to 32-cell

Unfertilized *Phallusia* eggs injected with Ens::3GFP (green). Confocal images of 16-cell stage embryo showing microtubule behavior during interphase and mitosis. B5.2 pair of blastomeres are at the bottom. Note that the aster becomes asymmetric in both bottom blastomeres. Time in min. Scale bar = 30µm.

### Movie S4. Unequal Cell Division 3. 32 to 64-cell

Unfertilized *Phallusia* eggs injected with Ens::3GFP (green) and also with Kif2::Venus (also appears green) to label the CAB. Confocal images of 32-cell stage embryo showing microtubule behavior during interphase and mitosis. B6.3 pair of blastomeres are at the bottom. The CAB can be seen centered between the two pairs of centrosomes. Note that the upper spindle pole in each B6.3 blastomere approaches the CAB and midline and that the asters nearest the CAB become progressively smaller. Time min:sec. Scale bar = 30µm.

### Movie S5. 3D rendering of microtubules during UCD

Unfertilized *Phallusia* eggs injected with Ens::3GFP (green). 3D-renedered confocal z-planes showing astral microtubule behavior merged with brightfiled images from an 8-cell stage embryo. Note the size and behavior of the asters nearest the CAB (bottom right blastomeres) versus those far from the CAB in the B4.1 blastomeres.

### Movie S6. CAB visualization

Unfertilized *Phallusia* eggs injected with PH::Tom (red) to label the plasma membrane and also with EB3::GFP (green) to label microtubules. Embryo at the 8-cell stage with the two CAB blastomeres at the bottom. Several z-planes from a confocal time-series rendered in 3D to better visualize the CAB protrusion. Note the CAB can be seen as a protrusion in both blastomeres (red).

### Movie S7. Kif2 protein leaves the CAB during mitosis

Unfertilized eggs injected with EB3::GFP and Kif2::Tom then imaged using a confocal microscope during the 8 to 16-cell stage. Bright-field images are overlayed with the EB3::GFP and Kif2::Tom fluorescence images. Kif2 protein can be seen to leave the CAB during mitosis. Scale bar = 30 µm. Time in sec.

### Movie S8. Microtubule polymerization around the CAB

Fast confocal imaging of a cortical z-plane containing the CAB in isolated B5.2 blastomeres. Microtubules labelled with Ens::3GFP and the CAB with Kif2::Tom. Note that microtubules polymerize around the CAB and also occasionally in the hole in the center of the CAB. Scale bar = 5µm. Time in sec.

### Movie S9. Microtubule growth around the CAB in an intact embryo

Confocal images of Ens::3GFP and Kif2::Ven fluorescence during interphase of a 32 cell stage embryo. Microtubules can be seen to grow around the CAB but they do not penetrate the CAB. Scale bar = 20µm. Time in sec.

### Movie S10. Nocodazole pipette to depolymerize microtubules at the 8-cell stage

Embryos at the 8-cell stage (interphase) were bathed in Cell Mask orange to label the mitotic spindle poles and the plasma membrane and during mitosis a nocodazole pipette was applied to the surface of one B4.1 blastomere. Note the exaggerated movement of the spindle pole nearest the nocodazole pipette towards the surface of the blastomere and CAB. Time in min:sec.

### Movie S11. Nocodazole pipette to depolymerise CAB proximal microtubules in zygote

Zygote with microtubules labelled with Ens::3GFP. A nocodazole pipette was advanced towards one aster and led to the loss of microtubules on one side of the zygote. Time in min:sec.

## References

1. Garzon-Coral, C., Fantana, H. A. & Howard, J. A force-generating machinery maintains the spindle at the cell center during mitosis. Science 352, 1124–1127 (2016).

2. Kern, D. M., Nicholls, P. K., Page, D. C. & Cheeseman, I. M. A mitotic SKAP isoform regulates spindle positioning at astral microtubule plus ends. J. Cell Biol. 213, 315–328 (2016).

3. Minc, N. & Piel, M. Predicting division plane position and orientation. Trends Cell Biol. 22, 193–200 (2012).

4. Cowan, C. R. & Hyman, A. A. ASYMMETRIC CELL DIVISION IN C. ELEGANS: Cortical Polarity and Spindle Positioning. Annu. Rev. Cell Dev. Biol. 20, 427–453 (2004).

5. Kotak, S. & Gönczy, P. Mechanisms of spindle positioning: cortical force generators in the limelight. Curr. Opin. Cell Biol. 25, 741–748 (2013).

6. Homma, N. et al. Kinesin superfamily protein 2A (KIF2A) functions in suppression of collateral branch extension. Cell 114, 229–239 (2003).

7. Maor-Nof, M. et al. Axonal pruning is actively regulated by the microtubule-destabilizing protein kinesin superfamily protein 2A. Cell Rep. 3, 971–977 (2013).

8. Siller, K. H. & Doe, C. Q. Spindle orientation during asymmetric cell division. Nat. Cell Biol. 11, 365–374 (2009).

9. Wessel, G. M. et al. The biology of the germ line in echinoderms. Mol. Reprod. Dev. 81, 679–711 (2014).

10. Galli, M. & van den Heuvel, S. Determination of the cleavage plane in early C. elegans embryos. Annu. Rev. Genet. 42, 389–411 (2008).

11. Grill, S. W., Gönczy, P., Stelzer, E. H. & Hyman, A. A. Polarity controls forces governing asymmetric spindle positioning in the Caenorhabditis elegans embryo. Nature 409, 630–633 (2001).

12. Labbé, J. C., Maddox, P. S., Salmon, E. D. & Goldstein, B. PAR proteins regulate microtubule dynamics at the cell cortex in C. elegans. Curr. Biol. CB 13, 707–714 (2003).

13. Kozlowski, C., Srayko, M. & Nedelec, F. Cortical Microtubule Contacts Position the Spindle in C. elegans Embryos. Cell 129, 499–510 (2007).

14. Minc, N., Burgess, D. & Chang, F. Influence of cell geometry on division-plane positioning. Cell 144, 414–426 (2011).

15. Laan, L. et al. Cortical dynein controls microtubule dynamics to generate pulling forces that position microtubule asters. Cell 148, 502–514 (2012).

16. Lawrence, C. J. et al. A standardized kinesin nomenclature. J. Cell Biol. 167, 19–22 (2004).

17. Wordeman, L., Wagenbach, M. & von Dassow, G. MCAK facilitates chromosome movement by promoting kinetochore microtubule turnover. J. Cell Biol. 179, 869–879 (2007).

18. O’Rourke, S. M., Christensen, S. N. & Bowerman, B. Caenorhabditis elegans EFA-6 limits microtubule growth at the cell cortex. Nat. Cell Biol. 12, 1235–1241 (2010).

19. Gönczy, P. Mechanisms of asymmetric cell division: flies and worms pave the way. Nat. Rev. Mol. Cell Biol. 9, 355–366 (2008).

20. Galli, M. et al. aPKC phosphorylates NuMA-related LIN-5 to position the mitotic spindle during asymmetric division. Nat. Cell Biol. 13, 1132–1138 (2011).

21. Gotta, M., Dong, Y., Peterson, Y. K., Lanier, S. M. & Ahringer, J. Asymmetrically distributed C. elegans homologs of AGS3/PINS control spindle position in the early embryo. Curr. Biol. CB 13, 1029–1037 (2003).

22. Couwenbergs, C. et al. Heterotrimeric G protein signaling functions with dynein to promote spindle positioning in C. elegans. J. Cell Biol. 179, 15–22 (2007).

23. Severson, A. F. & Bowerman, B. Myosin and the PAR proteins polarize microfilament-dependent forces that shape and position mitotic spindles in Caenorhabditis elegans. J. Cell Biol. 161, 21–26 (2003).

24. Collins, E. S., Balchand, S. K., Faraci, J. L., Wadsworth, P. & Lee, W.-L. Cell cycle-regulated cortical dynein/dynactin promotes symmetric cell division by differential pole motion in anaphase. Mol. Biol. Cell 23, 3380–3390 (2012).

25. Kiyomitsu, T. & Cheeseman, I. M. Cortical dynein and asymmetric membrane elongation coordinately position the spindle in anaphase. Cell 154, 391–402 (2013).

26. Nguyen-Ngoc, T., Afshar, K. & Gönczy, P. Coupling of cortical dynein and Gα proteins mediates spindle positioning in Caenorhabditis elegans. Nat. Cell Biol. 9, 1294–1302 (2007).

27. Rabinowitz, J. S. & Lambert, J. D. Spiralian quartet developmental potential is regulated by specific localization elements that mediate asymmetric RNA segregation. Development 137, 4039–4049 (2010).

28. Lambert, J. D. Developmental Patterns in Spiralian Embryos. Curr. Biol. 20, R72–R77 (2010).

29. Holy, J. & Schatten, G. Differential behavior of centrosomes in unequally dividing blastomeres during fourth cleavage of sea urchin embryos. J. Cell Sci. 98 (Pt 3), 423–431 (1991).

30. Schroeder, T. E. Fourth cleavage of sea urchin blastomeres: microtubule patterns and myosin localization in equal and unequal cell divisions. Dev. Biol. 124, 9–22 (1987).

31. Hibino, T., Nishikata, T. & Nishida, H. Centrosome-attracting body: a novel structure closely related to unequal cleavages in the ascidian embryo. Dev. Growth Differ. 40, 85–95 (1998).

32. Prodon, F. et al. Dual mechanism controls asymmetric spindle position in ascidian germ cell precursors. Dev. Camb. Engl. 137, 2011–2021 (2010).

33. Nishikata, T., Hibino, T. & Nishida, H. The centrosome-attracting body, microtubule system, and posterior egg cytoplasm are involved in positioning of cleavage planes in the ascidian embryo. Dev. Biol. 209, 72–85 (1999).

34. Patalano, S. et al. The aPKC-PAR-6-PAR-3 cell polarity complex localizes to the centrosome attracting body, a macroscopic cortical structure responsible for asymmetric divisions in the early ascidian embryo. J. Cell Sci. 119, 1592–1603 (2006).

35. McDougall, A. et al. Centrosomes and spindles in ascidian embryos and eggs. Methods Cell Biol. 129, 317–339 (2015).

36. Ren, X. & Weisblat, D. A. Asymmetrization of first cleavage by transient disassembly of one spindle pole aster in the leech Helobdella robusta. Dev. Biol. 292, 103–115 (2006).

37. Paix, A., Chenevert, J. & Sardet, C. Localization and anchorage of maternal mRNAs to cortical structures of ascidian eggs and embryos using high resolution in situ hybridization. Methods Mol. Biol. Clifton NJ 714, 49–70 (2011).

38. Hirokawa, N. & Takemura, R. Kinesin superfamily proteins and their various functions and dynamics. Exp. Cell Res. 301, 50–59 (2004).

39. Hirokawa, N. & Tanaka, Y. Kinesin superfamily proteins (KIFs): Various functions and their relevance for important phenomena in life and diseases. Exp. Cell Res. 334, 16–25 (2015).

40. Paix, A., Le Nguyen, P. N. & Sardet, C. Bi-polarized translation of ascidian maternal mRNA determinant pem-1 associated with regulators of the translation machinery on cortical Endoplasmic Reticulum (cER). Dev. Biol. 357, 211–226 (2011).

41. Noda, Y. et al. Phosphatidylinositol 4-phosphate 5-kinase alpha (PIPKα) regulates neuronal microtubule depolymerase kinesin, KIF2A and suppresses elongation of axon branches. Proc. Natl. Acad. Sci. U. S. A. 109, 1725–1730 (2012).

42. Adams, D. W. & Errington, J. Bacterial cell division: assembly, maintenance and disassembly of the Z ring. Nat. Rev. Microbiol. 7, 642–653 (2009).

43. Aldridge, B. B. et al. Asymmetry and aging of mycobacterial cells lead to variable growth and antibiotic susceptibility. Science 335, 100–104 (2012).

44. Carminati, J. L. & Stearns, T. Microtubules orient the mitotic spindle in yeast through dynein-dependent interactions with the cell cortex. J. Cell Biol. 138, 629–641 (1997).

45. Martindale, M. Q. & Henry, J. Q. Intracellular fate mapping in a basal metazoan, the ctenophore Mnemiopsis leidyi, reveals the origins of mesoderm and the existence of indeterminate cell lineages. Dev. Biol. 214, 243–257 (1999).

46. Pang, K. et al. Genomic insights into Wnt signaling in an early diverging metazoan, the ctenophore Mnemiopsis leidyi. EvoDevo 1, 10 (2010).

47. Carré, D., Djediat, C. & Sardet, C. Formation of a large Vasa-positive germ granule and its inheritance by germ cells in the enigmatic Chaetognaths. Dev. Camb. Engl. 129, 661–670 (2002).

48. Prulière, G., Cosson, J., Chevalier, S., Sardet, C. & Chenevert, J. Atypical protein kinase C controls sea urchin ciliogenesis. Mol. Biol. Cell 22, 2042–2053 (2011).

49. Nipper, R. W., Siller, K. H., Smith, N. R., Doe, C. Q. & Prehoda, K. E. Galphai generates multiple Pins activation states to link cortical polarity and spindle orientation in Drosophila neuroblasts. Proc. Natl. Acad. Sci. U. S. A. 104, 14306–14311 (2007).

50. Siller, K. H., Cabernard, C. & Doe, C. Q. The NuMA-related Mud protein binds Pins and regulates spindle orientation in Drosophila neuroblasts. Nat. Cell Biol. 8, 594–600 (2006).

51. Cabernard, C., Prehoda, K. E. & Doe, C. Q. A spindle-independent cleavage furrow positioning pathway. Nature 467, 91–94 (2010).

52. Redemann, S. et al. Membrane invaginations reveal cortical sites that pull on mitotic spindles in one-cell C. elegans embryos. PloS One 5, e12301 (2010).

53. Gönczy, P. et al. zyg-8, a Gene Required for Spindle Positioning in C. elegans, Encodes a Doublecortin-Related Kinase that Promotes Microtubule Assembly. Dev. Cell 1, 363–375 (2001).

54. McDougall, A., Lee, K. W.-M. & Dumollard, R. Microinjection and 4D fluorescence imaging in the eggs and embryos of the ascidian Phallusia mammillata. Methods Mol. Biol. Clifton NJ 1128, 175–185 (2014).

55. Ganem, N. J. & Compton, D. A. The KinI kinesin Kif2a is required for bipolar spindle assembly through a functional relationship with MCAK. J. Cell Biol. 166, 473–478 (2004).

56. Sardet, C. et al. Embryological methods in ascidians: the Villefranche-sur-Mer protocols. Methods Mol. Biol. Clifton NJ 770, 365–400 (2011).

